# α-Synuclein impairs mitochondrial function and alters cryptochrome regulation in the substantia nigra

**DOI:** 10.64898/2026.08.18.745090

**Authors:** Sinéad A. O’Sullivan, Adrian Kacperczyk-Perdyan, Ayse Ulusoy, Rita Pinto-Costa, Shirley SL Lee, Urszula Ławrynowicz, Jochen Prehn, Jakub Mieczkowski, Donato A. Di Monte

**Affiliations:** German Center for Neurodegenerative Diseases (DZNE), Bonn 53127, Germany; International Research Agenda 3P, Medical University of Gdańsk, Poland; Royal College of Surgeons (RCSI), Dublin, Ireland

## Abstract

Dopaminergic neurons in the substantia nigra pars compacta are key targets of α-synuclein pathology and neurodegeneration in Parkinson’s disease (PD). It is thought that pathological accumulation of α-synuclein significantly contributes to nigral neuronal dysfunction and ensuing neuronal demise. In this study, we further assessed this possibility and interrogated the role of α-synuclein burden in compromising neuronal function and altering physiological neuronal pathways. In particular, we focused on nigral mitochondrial impairment and disruption of circadian regulatory pathways triggered by sustained α-synuclein expression. Using an *in vivo* AAV-mediated model, we show that α-synuclein accumulation over a period of 12 weeks is associated with mitochondrial complex I and IV deficits and leads to dopaminergic cell loss. Proximity ligation assays revealed association of both total and phosphorylated α-synuclein with mitochondrial proteins at a time (between 4 and 12 weeks) that paralleled the development of mitochondrial dysfunction. Spatial transcriptomic analysis of the substantia nigra identified coordinated alterations in genes involved in mitochondrial, metabolic, and circadian pathways, including increased expression of circadian-associated genes such as *Nr1d1, Nr1d2, Cry2, Arntl2, and Csnk1e*. At the protein level, α-synuclein overexpression was associated with a differential shift in cryptochrome protein expression, characterized by reduced CRY1 and increased CRY2. Data provide evidence of a specific window of time during which sustained α-synuclein burden results in direct α-synuclein-mitochondria interactions and nigral mitochondrial damage. During the same time period, a specific remodeling of molecular clock components occurs, providing a potential new mechanism contributing to metabolic and mitochondrial dysregulations and, ultimately, neuronal injury and degeneration.

## Introduction

Parkinson’s disease (PD) is a progressive neurodegenerative disorder characterized by the selective loss of dopaminergic neurons in the substantia nigra pars compacta (SNpc) and accumulation of α-synuclein aggregates^1^. Although α-synuclein burden is a central feature of PD, the mechanisms by which it contributes to neuronal dysfunction and degeneration remain poorly understood. Mitochondrial dysfunction is a well-established feature of PD pathogenesis, with impairments in oxidative phosphorylation, particularly complex I deficiency, consistently observed in both patients and experimental models^2^. α-Synuclein has been reported to associate with mitochondrial membranes and impair mitochondrial function, however, the temporal relationship between α-synuclein accumulation, mitochondrial dysfunction, and neuronal degeneration in vivo remains unclear^3^. In addition to mitochondrial dysfunction, circadian disturbances are increasingly recognized as a prominent feature of PD. At the biological level, circadian rhythms coordinate whole-body functions that include sleep-wake cycles, hormone secretion, and metabolic processes. Disruptions in these functions are common in PD patients, and increasing evidence suggests that these disturbances may exacerbate both motor and non-motor symptoms and potentially influence disease progression ^4–7^. At the molecular/cellular level, circadian clocks involve distinct, highly regulated pathways that are governed by transcriptional-translational feedback loops. In particular, CLOCK/BMAL complexes drive expression of Period (Per) and Cryptochrome (Cry) genes, whose protein products form repressive complexes that regulate their own transcription^8^. Beyond maintaining circadian rhythms, circadian clock components regulate key aspects of cellular metabolism, including mitochondrial respiration, redox homeostasis, oxidative stress responses and ATP production. Circadian regulation of these processes optimises cellular energy utilisation across the day-night cycle, whereas disruption of clock function has been associated with impaired mitochondrial activity and increased oxidative stress^9^. Among the core clock components, the cryptochrome proteins CRY1 and CRY2 function as principal transcriptional repressors of the molecular clock and are critical regulators of cellular metabolism and mitochondrial homeostasis.

Very little is known about alterations of cellular circadian clocks in PD and, in particular, whether cryptochrome expression is altered within injured dopaminergic neurons and whether pathological α-synuclein accumulation affects neuronal clock components remain unclear. The relationship between α-synuclein burden, circadian rhythm alterations and mitochondrial dysfunction may be particularly relevant for nigral dopaminergic cells since these neurons are characterised by high energetic demands and are particularly vulnerable to mitochondrial dysfunction and oxidative damage^10^. Emerging evidence indicates that circadian dysregulation influences dopaminergic signaling^11^, neuronal excitability^12^, and metabolic homeostasis^13^, potentially increasing neuronal vulnerability for neurodegenerative processes^12^. Furthermore, alterations in clock gene expression and rhythmicity have been reported in PD patients, including altered expression of core clock genes such as BMAL1, PER, and REV-ERB genes^14,15^. In the context of α-synuclein pathology, emerging evidence suggests interactions between α-synuclein and circadian regulation. Experimental models have reported alterations in clock gene expression and behavioural rhythms following α-synuclein overexpression, and genetic variation in circadian genes has been associated with PD risk and symptom severity^16,17^. However, whether α-synuclein directly alters circadian regulatory pathways within nigral dopaminergic neurons, and how such changes relate to mitochondrial dysfunction, remains to be elucidated.

Here, we combine in vivo α-synuclein overexpression with spatial transcriptomics and protein-level analyses to investigate the relationship between α-synuclein pathology, mitochondrial dysfunction, and circadian regulation in the mouse substantia nigra. We demonstrate that α-synuclein progressively associates with mitochondrial proteins, coinciding with impaired oxidative phosphorylation and coordinated transcriptional changes in metabolic and circadian pathways, including differential regulation of cryptochrome proteins. Together, these findings identify selective remodeling of molecular clock components as a previously underappreciated feature of α-synuclein pathology in dopaminergic neurons and highlight circadian regulatory pathways as potential contributors to PD pathogenetic processes.

## Materials and methods

### Surgical procedures

Animal experiments were approved by the State Agency for Nature, Environment and Consumer Protection in North Rhine Westphalia, Germany. Experiments were conducted in C57BL/NRJ female mice between 12 and 14 weeks of age. Animals were housed within individually ventilated cages, in a specific-pathogen free facility, and kept on a 12-h light/dark cycle with ad libitum access to food and water. Mice were anaesthetized with isoflurane and surgery was performed using a stereotaxic frame (stoelting) and a 5ul Hamilton syringe fitted with a pulled glass capillary tube (outer diameter of 60-80um). Animals received a single 1.5ul injection of AAVs encoding for human αSyn (5x10^12^gc/ml), which were injected unilaterally into the right substantia nigra pars compacta at the following coordinates: 2.3mm posterior and -1.1mm lateral to bregma and - 4.1mm ventral relative to the dura, which was calculated according to mouse atlas of Paxinos and Franklin. AAVs were injected at a rate of 0.2ul/min and the needle was left in position for an additional 5 min after the infusion was completed before being slowly retracted. The skin incision was then closed with metal clips (Michel clips) and animals were allowed to recover in a heated recovery cage prior to being returned to their home cage.

### Viral vectors

Adeno-associated viral vectors (AAVs) used in this study were human α-synuclein or green fluorescent protein (GFP) and were generated using a backbone plasmid of AAV2-derived genome encapsulated into an AAV6 capsid. Expression of human α-synuclein or GFP were driven by the synapsin promoter. A woodchuck-hepatitis virus post-transcriptional regulatory element (WPRE) and a polyadenylation signal sequence were inserted downstream to the promoter and transgene sequence. Production and titration of the AAVs were carried out by Sirion Biotech. Stock titer AAV preparations were diluted in phosphate-buffered saline. Working titre was 5x10^12^ genome copies/ml.

### Immunohistochemistry

Immunohistochemical analysis of mitochondrial respiratory chain subunits and circadian proteins was performed on free-floating 30 μm coronal brain sections using fluorescence microscopy. All animals were transcardially perfused with 4% paraformaldehyde (PFA) within the same 4-hour time window to minimise potential circadian variation in protein expression. Brain tissue was collected at 4 and 12 weeks following AAV injection. Staining was carried out with free floating sections using the following primary antibodies: anti-tyrosine hydroxylase (TH, a marker of dopaminergic cells; Millipore ab152 and ab1542), anti-human α-synuclein (Sigma 36-008), anti-phosphorylated α-synuclein (WAKO 010-26481), anti-Grp75 (a mitochondrial marker; Abcam ab53098), anti-GRIM19 (a marker of mitochondrial complex I; Abcam ab110240) or anti-MTCO1 (a marker of complex IV; Abcam ab14705), anti-Cry1 (Proteintech 13474-1-AP), anti-Cry2 (Proteintech 13997-1-AP), anti-Bmal1 (Proteintech 14268-1-AP). Fab fragments were used to convert rabbit anti-Grp75 into donkey (Jackson Immuno 711-007-003). Secondary antibodies used were Goat anti-IgG2a mouse 488 (Thermofisher A21131), goat anti-donkey 594 (Abcam ab98822), goat anti-IgG1 mouse 647 (Invitrogen A21240) and goat anti-rabbit 405 (Jackson Immuno 111-475-003). Briefly, sections were washed in Tris-HCL followed by antigen retrieval with citrate buffer for 5 min at 95 °C. After washing with Tris-HCL, blocking/permeabilization was carried out with 5% normal goat serum and 0.5% Triton X-100 overnight at 4 °C. Sections were washed in TBS and the first primary antibodies (anti-Grim19 and anti-Grp75) were incubated with 1% BSA for 2 days at 4 °C. Following a wash step, sections were incubated with fab fragments overnight at 4 degrees. Another wash step preceeded the first fluorescent secondary antibodies (goat anti-mouse 488 and goat anti-donkey 594). Sections were then blocked with 5% normal mouse and 5% normal rabbit serum. The second primary antibodies (anti-alpha synuclein and TH) were incubated for 2 days at 4 °C. Wash steps with TBS were followed by the second fluorescent secondary antibodies (goat anti-mouse 647 and goat anti-rabbit 405). Sections were mounted onto coated glass slides and coverslipped using Vectashield hard set mounting media (H-1400).

### Image analysis

For quantitative image analysis of TH-positive neurons within the substantia nigra, two midbrain sections per animal were analysed. Images were acquired systematically across the substantia nigra from medial to lateral regions using Zeiss LSM880 or LSM980 confocal microscopes equipped with Airyscan detectors. Confocal *z*-stacks were acquired using ×20 or ×40 Plan-Apochromat objectives. Images were processed using ZEN software (Carl Zeiss) and analysed using Imaris 3D software (version 9.5; Oxford Instruments) or ImageJ.

For analysis of mitochondrial respiratory chain proteins within dopaminergic neurons, separate threshold-based surfaces were generated for TH, GRIM19 or MTCO1, and GRP75 immunoreactivity. TH-positive surfaces were used to define individual dopaminergic neurons as regions of interest. GRP75-positive signal was subsequently masked within each TH-positive neuron to identify the mitochondrial compartment, followed by masking of GRIM19 or MTCO1 signal within the GRP75-positive mitochondrial region. This approach enabled quantification of mitochondrial respiratory chain proteins specifically within individual TH-positive neurons. Raw fluorescence intensity values were obtained for GRIM19 or MTCO1, GRP75, and α-synuclein. GRIM19 and MTCO1 intensities were normalized to GRP75 to account for mitochondrial content, generating GRIM19:GRP75 and MTCO1:GRP75 ratios, respectively.

For quantification of CRY1, CRY2, and BMAL1 immunoreactivity within dopaminergic neurons, single-plane confocal images were analysed using ImageJ. Individual TH-positive neurons were manually delineated to generate cellular regions of interest (ROIs). CRY1, CRY2, or BMAL1 fluorescence was independently thresholded and quantified within the TH-positive ROIs. Depending on the analysis, percentage positive area and integrated fluorescence density were measured within the defined TH-positive regions.

For analysis of nuclear BMAL1, DAPI staining was used to generate nuclear masks in the image field of view. The BMAL1 channel was subsequently filtered through the DAPI-positive nuclear mask, and BMAL1 integrated density was quantified within the resulting nuclear ROIs.

For CRY1 spatial heterogeneity, the coefficient of variation (CV) of fluorescence intensity was calculated within individual TH-positive ROIs as the standard deviation of pixel intensity divided by the mean pixel intensity. Where multiple neurons were analysed from an individual animal, individual cellular measurements were averaged to generate a single value per animal.

### Western Blotting

Protein levels were measured using fresh-frozen substantia nigra tissue collected from PBS and AAV-injected mice. Both contralateral and ipsilateral substantia nigra were dissected and mechanically homogenized using a Precellys24 Touch Homogenizer (Bertin Technologies) in ice-cold lysis buffer consisting of RIPA buffer (Sigma-Aldrich) supplemented with 1× protease and phosphatase inhibitor cocktail (Thermo Fisher Scientific; Cat. No. 78429). Homogenates were centrifuged at 14,000 × g for 20 min at 4°C, and the supernatants were collected for protein analysis. Protein concentration was determined using the Pierce™ BCA Protein Assay Kit (Thermo Fisher Scientific), and 5 μg of total protein was loaded onto 4–12% Tris-glycine SDS-PAGE gels. Samples were either heated to 37°C or denatured at 95°C for 5 min in Laemmli sample buffer supplemented with 5% β-mercaptoethanol prior to electrophoresis. Proteins were transferred to PVDF membranes using a wet transfer system. Membranes were blocked in 5% bovine serum albumin (BSA) in PBS for 1 hr at room temperature before incubation with primary antibodies overnight at 4°C. Following three washes (10 min each) in PBS containing 0.05% Tween-20 (PBS-T), membranes were incubated with horseradish peroxidase (HRP)-conjugated goat anti-mouse (1:5,000; Cat. No. #1706516) or goat anti-rabbit (1:5,000; Cat. No. 1706515) secondary antibodies for 1 h at room temperature. Protein bands were visualized using an enhanced chemiluminescence (ECL) substrate (Cat. No. WBKLS0050) and imaged using a ChemiDoc™ Imaging System (Bio-Rad). Band intensities were quantified using ImageJ software (version 1.54f; RRID:SCR_003070).

### Proximity Ligation assay

Proximity ligation assays (PLA) were performed to assess the spatial proximity between α-synuclein and mitochondrial proteins in the substantia nigra using the Duolink Kit (Sigma-Aldrich), according to the manufacturer’s instructions. Briefly, mice were perfused and brains were fixed in 4% paraformaldehyde, cryoprotected with 30% sucrose, and sectioned at 35 µm. Free-floating sections containing the substantia nigra were permeabilized in PBS containing 0.25% Triton X-100 and blocked using Duolink blocking solution for 1 hour at 37°C. Sections were incubated overnight at 4°C with combinations of primary antibodies, including antibodies against α-synuclein (Syn211), phosphorylated α-synuclein (pS129), and mitochondrial proteins TOM20, GRP75, and LONP1. The following antibody pairs were used: TOM20/Syn211, TOM20/pS129, GRP75/Syn211, GRP75/pS129, and LONP1/Syn211. After washing, sections were incubated with species-specific PLA probes (PLUS and MINUS) for 1 hour at 37°C. Ligation and rolling circle amplification were performed according to the manufacturer’s protocol, resulting in the formation of discrete PLA puncta indicating protein proximity (<40 nm). Sections were mounted and imaged using brightfield microscopy. PLA signal (puncta) was quantified within the substantia nigra pars compacta (SNpc).

### Stereological counting

Analyses were performed on midbrain sections containing the substantia nigra, by an investigator blinded to the sample IDs. Unbiased stereological estimates of the number of nigral neurons were obtained by counting under brightfield microscopy. Samplings were performed on every fifth section throughout the entire SNpc. Delineations were made using a 4x objective, and counting was performed using a 63x Plan-Apo oil objective (Numerical aperture = 1.4). A guard zone thickness of 1 μm was set at the top and bottom of each section. Cells were counted using the optical fractionator technique (Stereo Investigator software version 9, MBF Biosciences, Williston, VTA, USA) using a motorized Olympus microscope (IX2 UCB) equipped with an Olympus disk spinning unit (DSU) and a light sensitive EM-CCD camera. Coefficient of error was calculated according to (Gundersen and Jensen, 1987); values < 0.10 were accepted.

### Complex I Dipstick Assay

Complex I activity was measured using an immunocapture-based dipstick assay (ab109720, Abcam). Dipsticks containing immobilized Complex I were immersed in a reaction solution containing NADH and nitro blue tetrazolium (NBT). Active Complex-I oxidizes NADH, leading to the reduction of NBT and formation of a bluish-purple precipitate at the antibody capture line. Fresh tissue samples were rapidly dissected and homogenized in extraction buffer according to the manufacturer’s instructions. Protein concentration was determined, and sample volume was adjusted to 25 µL containing 10 µg of total protein prior to incubation with the dipsticks.

### Spatial Transcriptomics

Mice were euthanized 12 weeks following unilateral injection of AAV-human α-synuclein or PBS into the midbrain containing the substantia nigra. Brains were collected, fixed in 4% paraformaldehyde (PFA), and paraffin embedded.

### FFPE RNA quality assessment

RNA quality was evaluated by extracting total RNA from representative FFPE tissue sections using the RNeasy FFPE Kit (Qiagen). RNA integrity was assessed using the DV200 metric, defined as the percentage of RNA fragments longer than 200 nucleotides. Only FFPE blocks with DV200 ≥ 60% were included in the spatial transcriptomic analysis.

### FFPE tissue sectioning

FFPE blocks were sectioned at the Department of Pathomorphology, Medical University of Gdansk. Tissue sections were mounted onto standard glass slides compatible with the Visium CytAssist workflow. The slides were incubated for 3 h at 42°C and subsequently dried overnight at room temperature. Before further processing, the tissue sections and selected regions of interest were verified to fit within the 6.5-mm Visium CytAssist capture area.

### H&E staining, brightfield imaging, and tissue preprocessing

Tissue sections were deparaffinized, stained with hematoxylin and eosin, coverslipped, and imaged in brightfield mode according to the Visium CytAssist Spatial Gene Expression for FFPE deparaffinization, H&E staining, imaging, and decrosslinking protocol (10x Genomics, CG000520). Brightfield images were acquired to document tissue morphology, confirm tissue integrity, and enable subsequent registration of the spatial gene expression data. After imaging, the coverslips were removed, and the sections were destained and decrosslinked. The processed tissue slides were subsequently subjected to overnight hybridization with mouse whole-transcriptome probe pairs at 50°C according to the Visium CytAssist Spatial Gene Expression Reagent Kits User Guide (10x Genomics, CG000495).

### Visium CytAssist library construction and sequencing

Following probe hybridization, unbound probes were removed and adjacent probe pairs hybridized to their target transcripts were ligated. The tissue slides and 6.5-mm Visium CytAssist Spatial Gene Expression slides were loaded into the Visium CytAssist instrument, which mediated the release and spatially registered transfer of the ligated probe products from the tissue sections onto the barcoded capture areas. Probe extension, elution, pre-amplification, and library construction were performed using the Visium CytAssist Spatial Gene Expression for FFPE, Mouse Transcriptome, 6.5-mm reagent kit (10x Genomics, PN-1000521), following the manufacturer’s instructions (CG000495).

The number of cycles used for sample-index PCR was determined by qPCR using a LightCycler 480 instrument. The measured Cq values ranged from 9.2 to 9.8, and the final number of amplification cycles was set to Cq + 2. Libraries were indexed using the 10x Genomics Dual Index Kit TS Set A (PN-1000251) and quantified by qPCR using the KAPA Library Quantification Kit. The target sequencing depth was calculated as the fraction of the 6.5-mm capture area covered by tissue multiplied by 4,992 capture spots and 25,000 read pairs per tissue-covered spot. Pooled libraries were sequenced on an Illumina NextSeq 550 using the following paired-end configuration: Read 1, 28 cycles; i7 index, 10 cycles; i5 index, 10 cycles; and Read 2, 50 cycles. The Dual Index Kit TS Set A is required for this assay, and the NextSeq 500/550 platform and stated read configuration are compatible with Visium CytAssist libraries.

### VSGE data processing

Raw sequencing data (BCL files) were converted to FASTQ format using Space Ranger (v2.0.0). Reads were aligned to the mouse reference genome (mm10) using STAR, and spatial feature counts were quantified with Visium Mouse Transcriptome Probe Set v1.0. Loupe Browser 8.0 was used for alignment visualization and quality control.

### Data preprocessing, clustering, and differential expression

Spatial transcriptomic data were processed in Seurat (v5.3) using the *Load10X_Spatial* function. Low-quality spots were excluded based on feature counts (<100 or >8000) or mitochondrial gene content (>15%). Data were normalized using *NormalizeData* and *SCTransform* functions with default parameters. Highly variable features were identified using variance-stabilizing transformation (VST), and data were scaled with regression of mitochondrial gene content. Dimensionality reduction was performed using PCA followed by UMAP. Nearest neighbours were computed using the first 20 principal components based on ElbowPlot, and clustering was performed using a shared nearest-neighbour (SNN) algorithm at a resolution of 0.8. Differentially expressed genes (DEGs) were identified using *FindAllMarkers* (only.pos = TRUE, test.use = “MAST”, min.pct = 0.25, logfc.threshold = 0.25), retaining genes with P < 0.05. MAST is a flexible framework for single-cell RNA-seq that models both the probability of expression and the expression level, enabling supervised differential expression analyses and unsupervised exploration of co-expression patterns. Seurat’s Cell Selector was used to align tissue regions of interest.

### Cell type deconvolution

Robust Cell Type Decomposition (RCTD) was performed using a single-cell RNA reference deposited in GEO under accession code GSE153424 (samples GSM55, GSM57, GSM68). Single-cell data were filtered based on gene count quantiles (2.5–97.5%) and mitochondrial content (<5%), clustered at resolution 0.5, and annotated using canonical marker genes. RCTD was run in doublet mode, and deconvoluted cell-type proportions were visualized using *SpatialFeaturePlot*.

### Statistical analysis

Data are presented as mean ± standard error of the mean (SEM), unless otherwise stated. Statistical analyses were performed using GraphPad Prism (version 10; GraphPad Software, Boston, MA, USA). Comparisons between two groups were performed using two-tailed Student’s *t*-tests. Where ipsilateral and contralateral regions from the same animal were compared, paired *t*-tests were used; comparisons between independent experimental groups were performed using unpaired *t*-tests. Comparisons involving more than two groups were performed using one-way analysis of variance (ANOVA), followed by the appropriate multiple-comparisons post hoc test as specified in the corresponding figure legends. Correlations were assessed using Spearman’s correlation analysis. Statistical significance was defined as *P* < 0.05. Individual animals were considered biological replicates.

## Results

### Human α-synuclein overexpression in the substantia nigra induces dopaminergic neuron loss over time

To determine the impact of α-synuclein-mediated pathology on dopaminergic neuron survival, AAVs encoding human α-synuclein were unilaterally injected into the substantia nigra of adult female C57BL/6 mice (Fig. 1A). Robust human α-synuclein protein expression was observed in ventral mesencephalic tissue containing the substantia nigra. Quantification by western blot at 4 and 12 weeks post-injection revealed a significant reduction at the later time point (Fig. 1B). Stereological analysis revealed that a marked, significant reduction of dopaminergic neurons occurred in the injected substantia nigra pars compacta (SNpc) between the 4- and 12-week time points; this loss was indicated by counting TH-immunoreactive and Nissl-positive nigral neurons (Fig. 1C–E). In contrast, control AAV-GFP injection did not result in significant dopaminergic neuron loss, indicating that the neuronal loss observed following α-synuclein overexpression was not attributable to AAV delivery alone (Fig. 1C-D). Human α-synuclein protein expression was also quantified at the cellular level by densitometric analysis of human α-synuclein immunofluorescence (Syn211) within nigral TH-positive neurons. Interestingly, no difference was observed between measurements at 4 and 12 weeks post-injection (Fig. 1F–G). These data are apparently at odds with the results described above showing a reduction of human α-synuclein protein in whole tissue samples (please compare Fig. 1F–G *vs.* Fig. 1B). However, taken together, these findings indicate that α-synuclein overexpression is decreased over time in tissue samples as a result of nigral neuronal loss; it instead remains relatively unchanged when assessed within individual dopaminergic neurons at either the 4- and 12-week time points. A recognised feature of pathological α-synuclein is its phosphorylation at the serine 129 (pS129) residue. In our model, phosphorylated α-synuclein levels were significantly increased at 12 weeks compared with 4 weeks (Fig. 1H–I), consistent with progressive accumulation of pathological α-synuclein species even in the absence of significant changes in total human α-synuclein expression. Therefore, although the α-synuclein burden within TH-positive neurons remained relatively stable over time, this was accompanied by increased α-synuclein phosphorylation and greater dopaminergic neuron loss at 12 weeks. These findings suggest that the development of neuronal pathology is not simply associated with increasing total α-synuclein abundance, but rather with time-dependent changes in the pathological state of α-synuclein and/or downstream cellular responses to its sustained expression.

**Figure 1:**
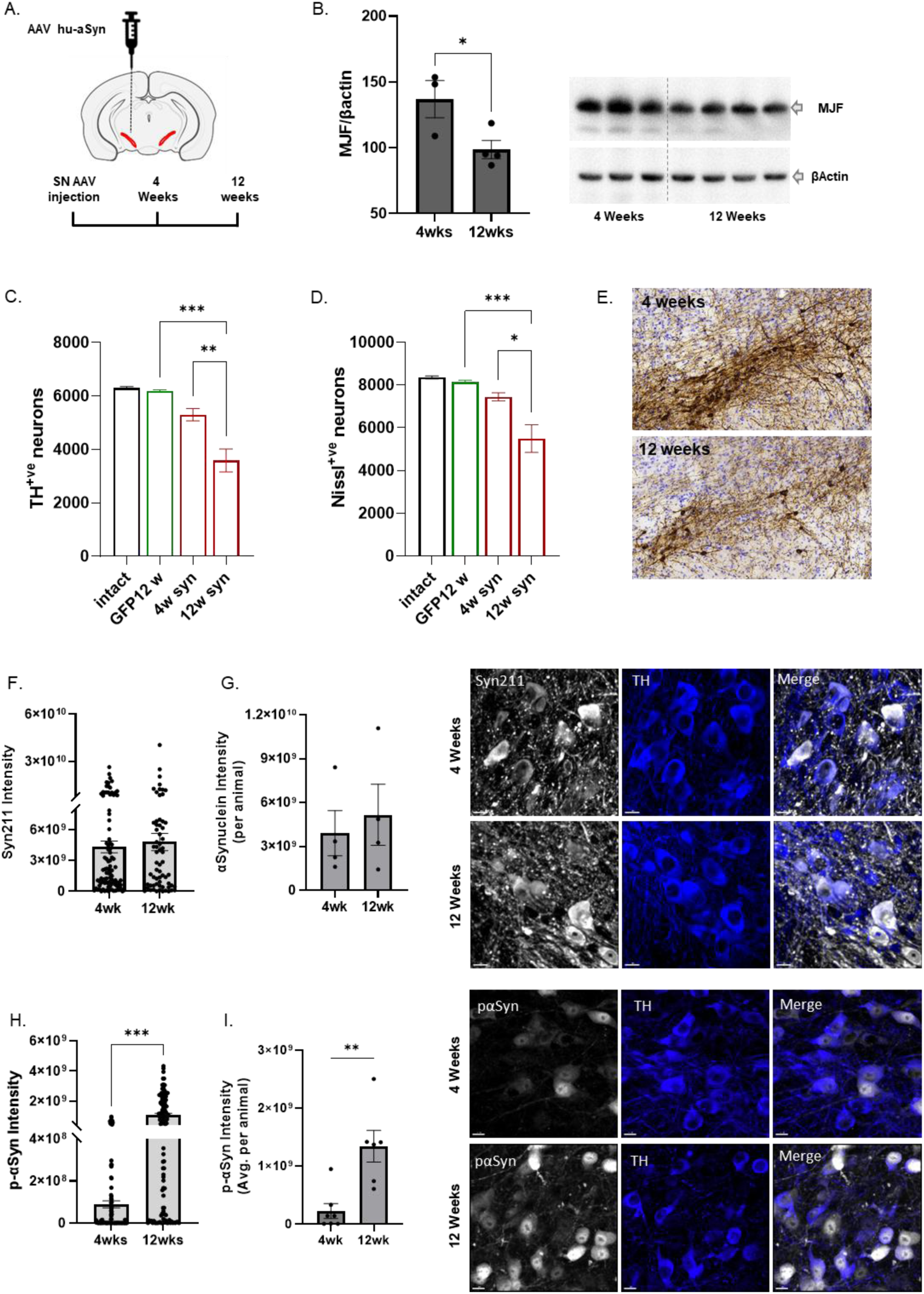
Human α-synuclein overexpression in the substantia nigra induces dopaminergic neuron loss over time. A. Schematic illustrating unilateral injection of AAV human αSyn at 5x10^12^ gc/ml into the SN of C57BL/6 mice. B. Western blot showing total α-synuclein in the SN at 4- and 12-weeks post-injection. C,D. stereological quantification of TH-positive neurons counterstained with Nissl E. Representative images of SN tissue sections used for stereology stained for TH and Nissl. F,G Quantification and representative images of total α-synuclein (Syn211) immunofluorescence within TH-positive neurons. H,I. Quantification and representative images of immunofluorescent staining for phosphorylated αSyn (pS129) within TH positive neurons. Scale bar 10µm. N=5-6 mice per group, statistics used were student’s t-test and one-way ANOVA. *p<0.05, **p<0.01, ***p<0.001

### α-Synuclein overexpression induces mitochondrial complex I and IV deficits in dopaminergic neurons

Given the central role of mitochondrial dysfunction in Parkinson’s disease, we next investigated whether α-synuclein overexpression disrupts the expression of mitochondrial oxidative phosphorylation (OXPHOS) proteins in dopaminergic neurons and the time course of this dysfunction. Mitochondrial OXPHOS components were quantified within TH-positive neurons of the substantia nigra. Analyses focused on complexes I and IV, both of which are critical components of the mitochondrial respiratory chain and have previously been reported to be reduced in postmortem substantia nigra tissue from individuals with PD^18^. At 4 weeks post-injection, no significant changes in complex I (as assessed by quantification of one of its subunits, GRIM19) or complex IV (as assessed by quantification of its MTCO1 subunit) levels were observed (Fig. 2A, B and 2E, F). In contrast, by 12 weeks, α-synuclein overexpression resulted in a significant reduction in both neuronal complex I and complex IV (Fig. 2C, D and 2G, H respectively and supplemental Fig 1A-D). To determine whether mitochondrial deficits were associated with intraneuronal α-synuclein burden, we performed a correlation analysis between complex I levels and total α-synuclein (Syn211) intensity. No significant relationship was observed at 4 weeks (Fig. 2I); however, a significant negative correlation emerged at 12 weeks (Fig. 2J) indicating that, at this later time point, greater α-synuclein burden was associated with reduced Complex I levels. To assess functional consequences, we measured complex I enzymatic activity using a dipstick assay. Consistent with protein-level changes, complex I activity was significantly reduced at 12 weeks but not at 4 weeks (Fig. 2K). To control for potential effects of AAV-mediated transgene expression itself, mitochondrial protein levels were also assessed in mice injected with AAV encoding GFP. No corresponding mitochondrial deficits were observed in AAV-GFP–injected mice, supporting an α-synuclein-specific effect rather than a general consequence of viral transduction or transgene overexpression (Supplementary Fig. 2A–B and Fig. 1C). Together, these findings demonstrate that loss of complex I and complex IV levels within nigral dopaminergic neurons primarily occur between 4 and 12 weeks post AAV injection, paralleling the accumulation of pathological, phosphorylated α-synuclein and the development of neurodegeneration.

**Figure 2:**
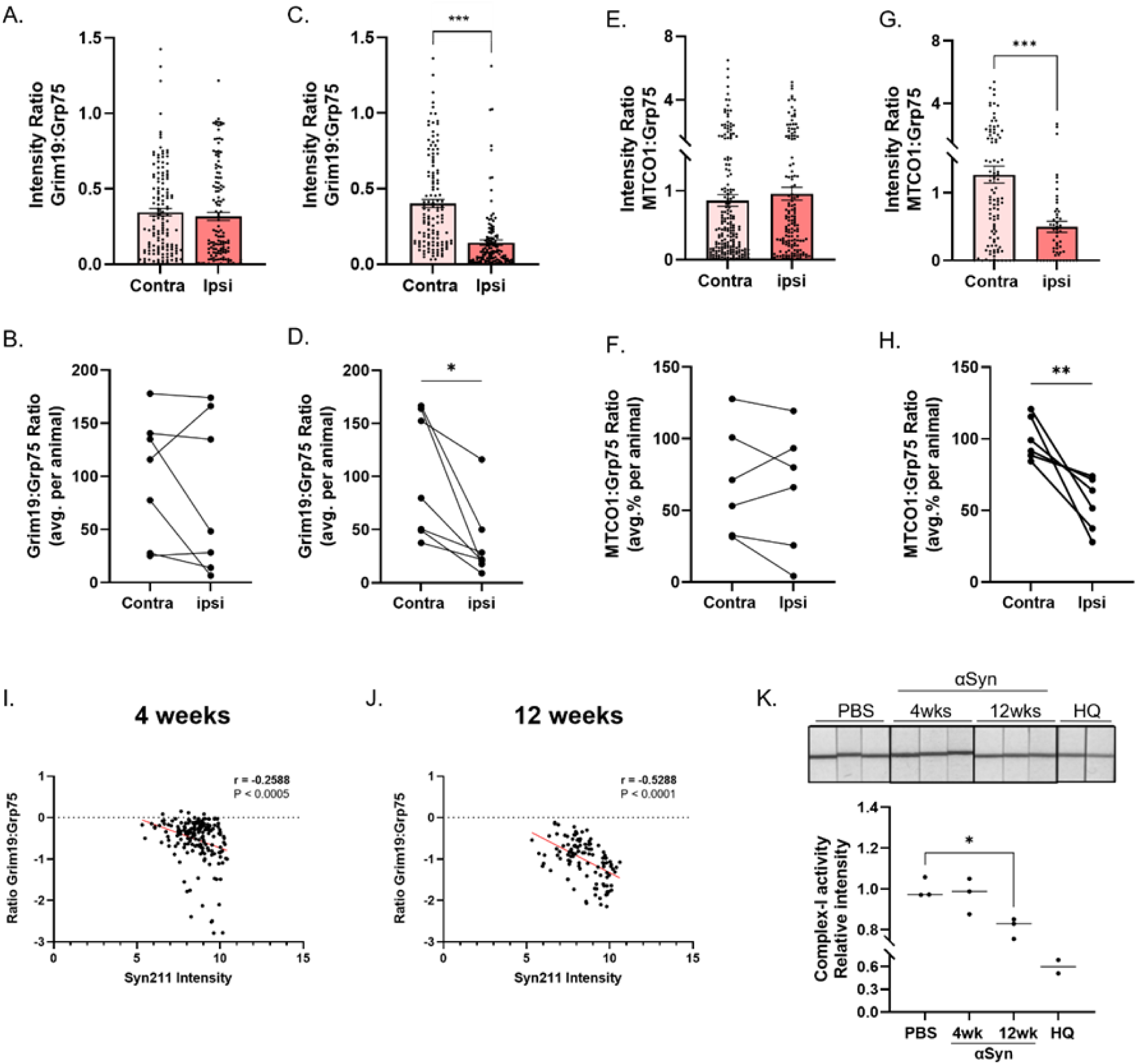
α-Synuclein overexpression induces mitochondrial complex I and IV deficits in dopaminergic neurons. A–D. Quantification of Complex I (GRIM19:GRP75) in TH-positive neurons at 4 and 12 weeks following AAV-human α-synuclein injection, shown as per-cell and per-animal measurements. E–H. Corresponding quantification of Complex IV (MTCO1:GRP75). I-J. Correlation between cellular GRIM19:GRP75 intensity ratio and Syn211 immunofluorescence intensity at 4 and 12 weeks. K. Representative Complex I dipstick assay and quantification of Complex I activity in the substantia nigra from PBS, α-synuclein, and Harlequin (HQ) mice. N=3-7 mice per group, statistics used were student’s t-test and one-way ANOVA. *p<0.05, **p<0.01, ***p<0.001.

### α-Synuclein association with mitochondrial proteins increases over time in the substantia nigra

To determine whether mitochondrial dysfunction developing between 4 and 12 weeks was associated with time-dependent changes in the proximity of α-synuclein to mitochondrial proteins, we next assessed the spatial relationship between α-synuclein and mitochondrial proteins in vivo using proximity ligation assays (PLA). At 4 weeks post-injection, PLA signal assessing proximity between α-synuclein and the mitochondrial outer membrane protein TOM20 were detectable, albeit quite scarce; this α-synuclein-mitochondria association, however, became significantly more pronounced at 12 weeks (Fig. 3A, B). Similarly, total α-synuclein (Syn211) showed significant proximity with GRP75, another general mitochondrial marker, at the 12 week time point (Supplemental Fig. 3A). Given the accumulation of phosphorylated α-synuclein (pS129) observed in this AAV model (see Fig. 1), we next examined its association with mitochondrial proteins. PLA analysis revealed increased proximity between pS129 α-synuclein and either TOM20 or GRP75 at 12 weeks compared with 4 weeks (Fig. 3C and D respectively). Notably, PLA signal was also detected between α-synuclein and the mitochondrial matrix protein LONP1 (Supplementary Fig. 3B), with a significant increase at 12 weeks. Given the localization of LONP1 within the mitochondrial matrix, this finding is consistent with previous reports suggesting that α-synuclein may localize within mitochondria^19,20^, although PLA alone cannot establish mitochondrial import or intramitochondrial localization. Together, these data demonstrate a significant association of α-synuclein with mitochondrial proteins after sustained protein overexpression, temporally coinciding with the emergence of mitochondrial dysfunction.

**Figure 3:**
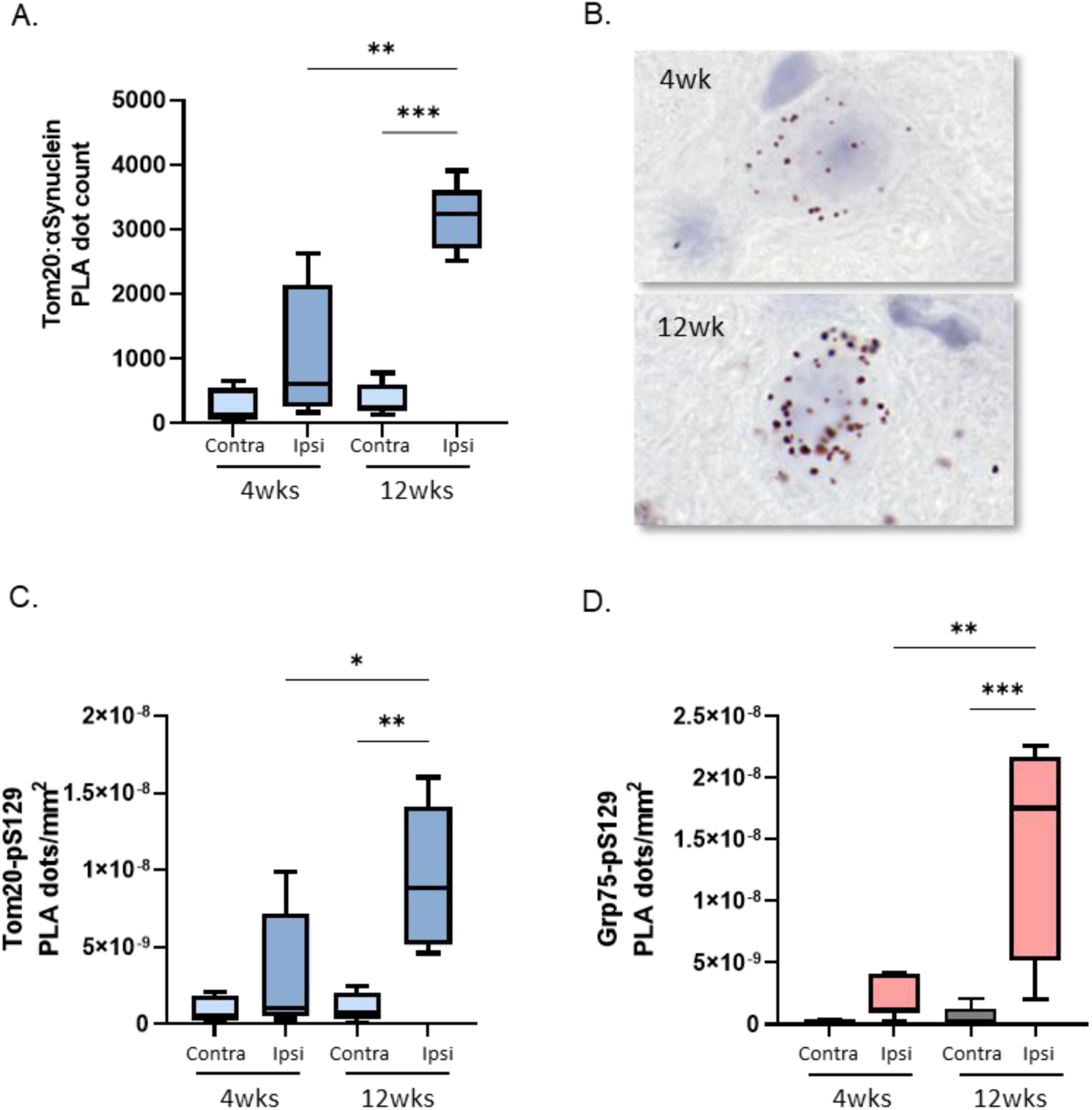
α-Synuclein association with mitochondrial proteins increases over time in the substantia nigra. Positive PLA puncta were quantified within the substantia nigra pars compacta (SNpc). A,B. Quantification and representative images of TOM20–Syn211 PLA signal at 4 and 12 weeks following α-synuclein overexpression. Graphs show quantification of PLA in contra- and ipsilateral SNpc of mice overexpressing human αSyn. C-D. Quantification of TOM20–pS129, GRP75–pS129 PLA signal. N=4-5 mice per group, statistics used were student’s t-test and one-way ANOVA. *p<0.05, **p<0.01.

### α-Synuclein overexpression alters mitochondrial and circadian gene expression in the substantia nigra

To investigate molecular pathways associated with α-synuclein-induced mitochondrial dysfunction, we performed Visium spatial transcriptomics on midbrain sections at 12 weeks post α-synuclein overexpression. SNpc-containing TH-enriched regions were selected for downstream analysis. Consistent with other α-synuclein models of PD^21^, differential gene expression analysis revealed significant alterations in genes associated with mitochondrial function and cellular metabolism (Fig. 4B). Decreased expression of complex I and complex IV subunits (e.g., *Ndufa9, Ndufb10, Cox5b*, and *Cox6c*) was also consistent with our mitochondrial deficits observed at the protein and functional levels (Fig. 2). Gene ontology analysis further identified enrichment of pathways related to metabolism and mitochondrial function. Quite interestingly it also revealed significant changes in circadian regulation pathways (Fig. 4C–D). To further interrogate these pathways, heatmaps of selected differentially expressed genes involved in mitochondrial function and circadian regulation were generated (Fig. 4E–F). Several circadian regulatory genes were upregulated, including *Nr1d1*, *Nr1d2*, *Csnk1d*, and *Csnk1e*, which are involved in transcriptional repression and post-translational regulation of the circadian clock^22^. Expression of the closely related cryptochrome genes differed, with Cry2 upregulated and Cry1 downregulated. As these paralogous genes are typically co-regulated within the molecular clock, this divergent expression pattern suggests altered regulation of the cryptochrome arm of the molecular clock. In addition, altered expression of other core clock genes, including *Per3* and *Arntl2* (*Bmal2*), further supports disruption of circadian regulatory pathways. Together, these findings indicate that α-synuclein overexpression is associated with coordinated transcriptional changes in mitochondrial and circadian regulatory pathways in the substantia nigra.

**Figure 4:**
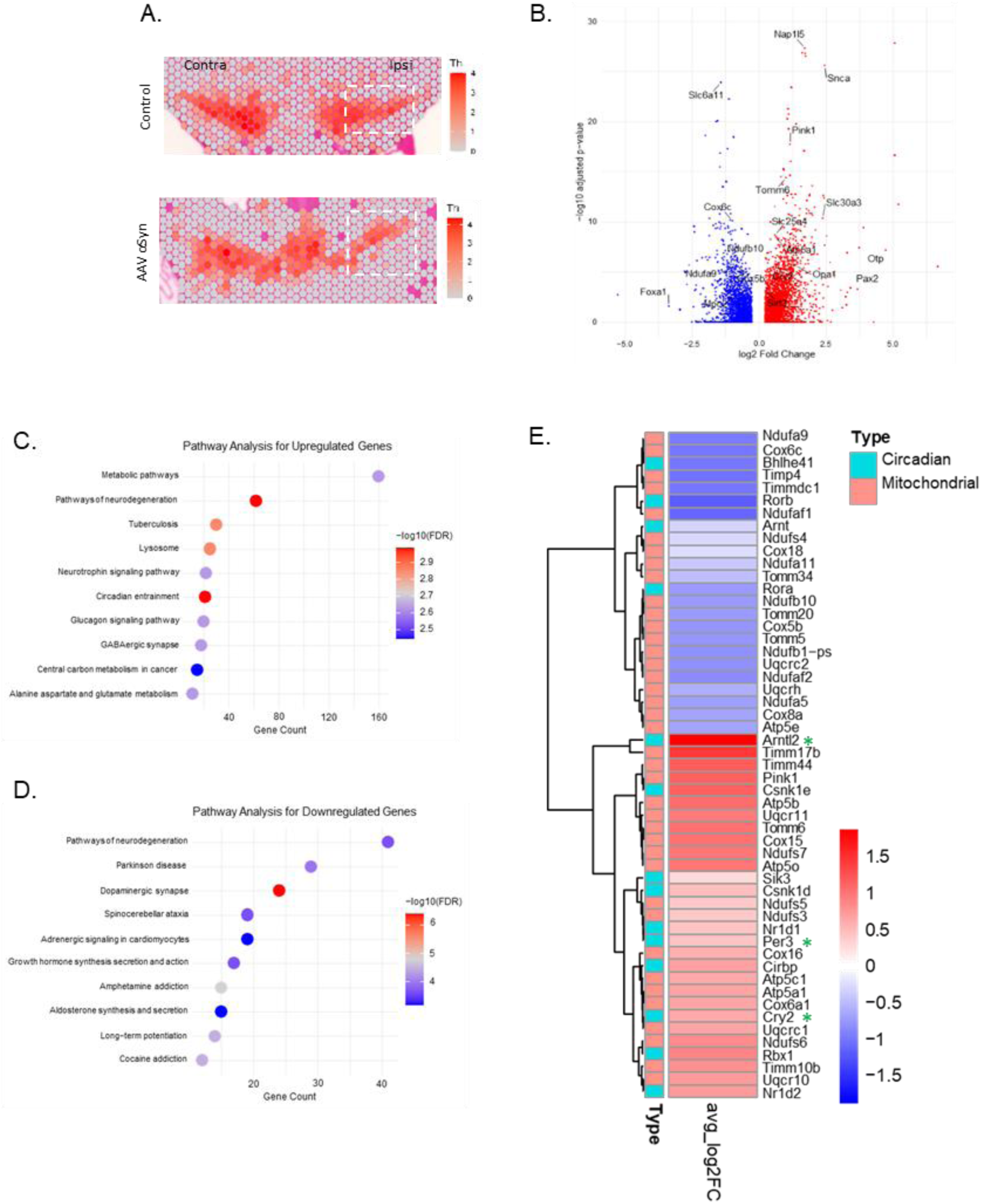
α-Synuclein overexpression alters mitochondrial and circadian gene expression in the substantia nigra. A. Spatial transcriptomic section showing the TH-enriched region of the substantia nigra used to guide selection for downstream analysis. Dashed outlines indicate the approximate region selected for analysis. B. Volcano plot of significantly up- and downregulated genes within selected ipsilateral substantia nigra regions following α-synuclein overexpression compared with PBS controls. C,D. Gene Ontology (GO) enrichment analysis of significantly upregulated and downregulated genes, respectively, performed using ShinyGO v0.80. **E.** Heatmap of selected differentially expressed genes associated with mitochondrial function and circadian rhythm. Red and blue indicate relative higher and lower expression, respectively. Cry2, Per3, and Arntl2 are highlighted as core circadian regulatory genes.

### α-Synuclein overexpression is associated with differential regulation of cryptochrome proteins in the substantia nigra

Given the observed changes in circadian gene expression, we next examined whether the core circadian repressors CRY1 and CRY2 were altered at the protein level. Immunofluorescent analysis revealed a significant increase in CRY2 and a reduction of CRY1 at 12 weeks as compared with GFP controls (Fig. 5A-E). The data with CRY1, albeit showing a reduction, did not reach statistical significance when expressed as integrated density, most likely due to the observed clustering of the fluorescent signal (Fig. 5C). This clustering was confirmed by analysis of CRY1 signal within TH-positive cells using the coefficient of variation; data showed a significantly higher coefficient of variation following α-synuclein overexpression, indicating a more heterogeneous intraneuronal distribution (Fig.5E). Final analyses therefore were carried out using the CRY1 percentage area; results confirmed a significant reduction of CRY1 at 12 weeks post AAV injection (Fig.5F). Together, these findings demonstrate differential cryptochrome protein regulation following α-synuclein overexpression, characterised by increased CRY2 and reduced CRY1 expression.

**Figure 5:**
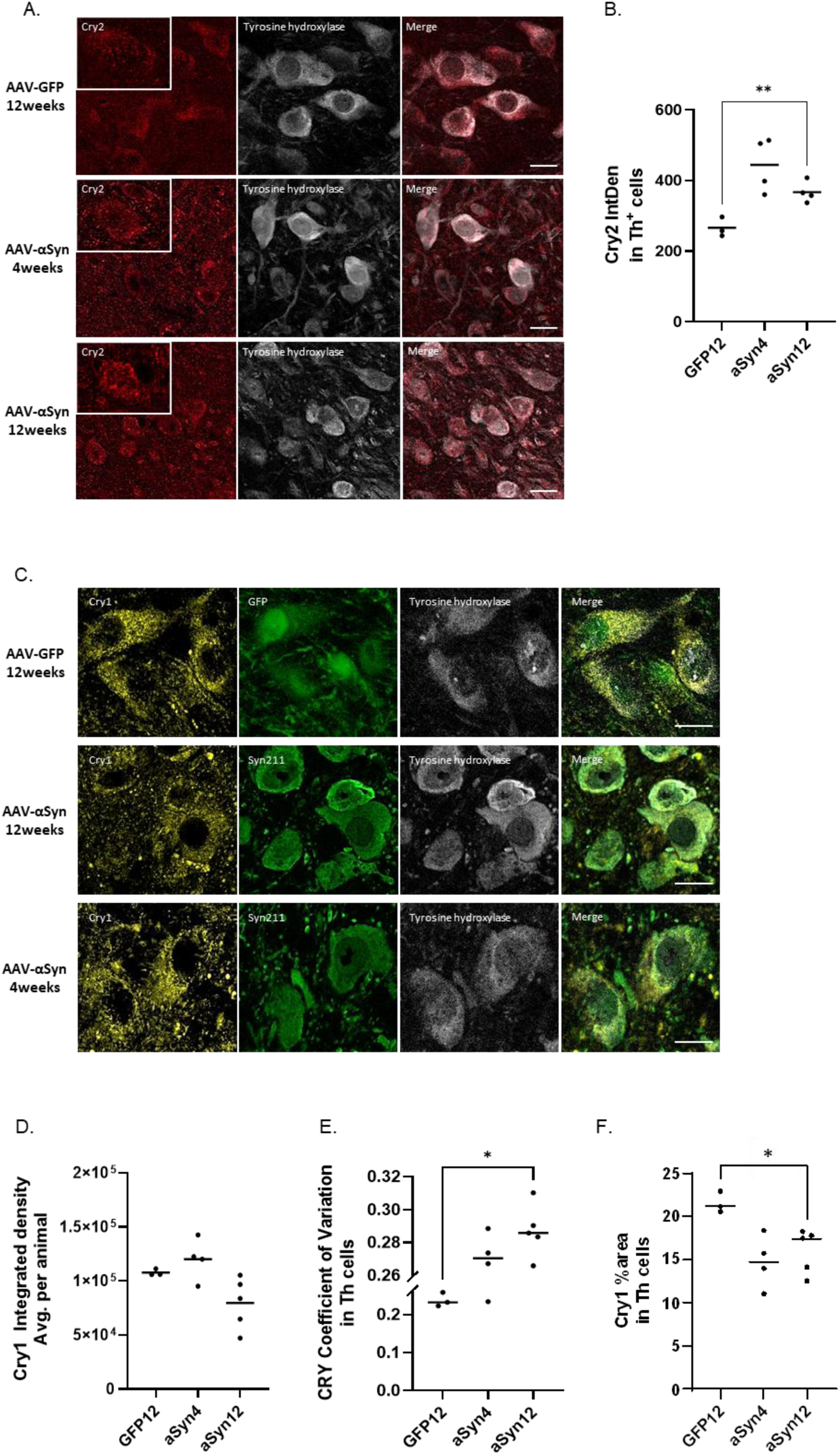
α-Synuclein overexpression is associated with differential cryptochrome expression in the substantia nigra. A. Immunofluorescent staining of Cry2 at 12 weeks following aSyn overexpression, compared with GFP controls. B. Quantification of Cry2 integrated density in TH-positive neurons. C. Representative Cry1 immunofluorescence with quantification of integrated density (D), coefficient of variation (E), and stained area (F). Graphs show mean ± SEM. Scale bar 10µm. One-way ANOVA with Dunnett’s post hoc test. *P<0.05, **P<0.01.

### BMAL1 protein expression is similar to GFP controls despite differential cryptochrome expression following α-synuclein overexpression

Given the differential regulation of the core circadian repressors CRY1 and CRY2 (Fig. 5), together with the increased expression of *Arntl2* (*Bmal2*) identified by spatial transcriptomics (Fig. 4), we investigated whether the positive transcriptional arm of the molecular circadian clock was similarly affected at the protein level. BMAL1 (ARNTL) is a core transcriptional activator that heterodimerizes with CLOCK to drive expression of *Per* and *Cry* genes, thereby functioning in opposition to cryptochrome-mediated transcriptional repression. Immunofluorescence analysis demonstrated no significant difference in nuclear BMAL1 integrated density between α-synuclein-overexpressing and GFP control mice at 12 weeks (Fig. 6A–B). Quantification of BMAL1 integrated density specifically within TH-positive neurons likewise revealed no significant difference between groups at 12 weeks (Fig. 6C). Together, these findings indicate that, unlike the reciprocal alterations observed in CRY1 and CRY2, BMAL1 protein expression was similar between groups at the 12-week endpoint following α-synuclein overexpression. This suggests that, in our AAV model, α-synuclein overexpression is associated with selective remodelling of cryptochrome proteins rather than broad changes in BMAL1 protein abundance at this time point.

**Figure 6:**
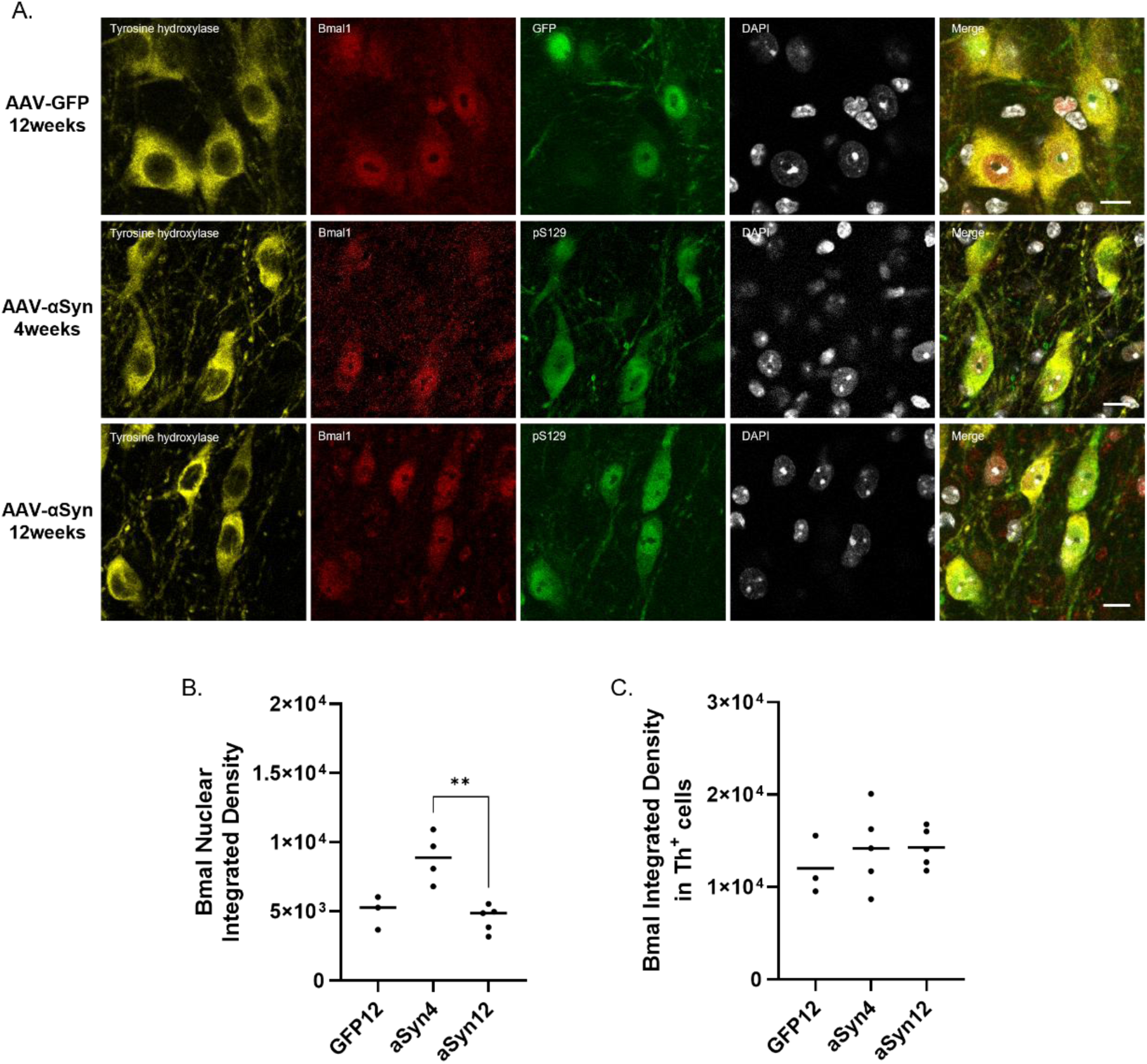
BMAL1 protein expression is similar to GFP controls despite differential cryptochrome expression following α-synuclein overexpression. A. Immunofluorescent staining of Bmal1 at 12 weeks following αSyn overexpression, compared with GFP controls. B. Quantification of nuclear BMAL1 integrated density. C. Quantification of BMAL1 integrated density within TH-positive neurons. Graphs show mean +/- SEM. One-way ANOVA with Dunnett’s post hoc test. Scale bar 10µm **P<0.01 for the indicated comparison.

## Discussion

In this study, we demonstrate that overexpression of human α-synuclein in the substantia nigra induces dopaminergic neuron loss over time accompanied by mitochondrial dysfunction and alterations in circadian regulatory pathways. Specifically, we show that sustained α-synuclein expression in dopaminergic neurons is accompanied by increasing α-synuclein-mitochondrial proximity over time and reductions in mitochondrial Complex I and Complex IV protein levels together with decreased Complex I activity. In parallel, spatial transcriptomic and protein analyses revealed alterations in circadian regulatory components, including differential expression of the cryptochrome proteins CRY1 and CRY2. Together, these findings suggest that α-synuclein pathology is associated with parallel alterations in mitochondrial function and circadian regulation within the substantia nigra.

Mitochondrial dysfunction is a well-established feature of Parkinson’s disease and α-synuclein toxicity. Our data extend previous observations by defining the temporal development of mitochondrial impairment in dopaminergic neurons following α-synuclein overexpression. At 4 weeks, despite robust α-synuclein expression, mitochondrial Complex I and IV levels and Complex I activity were largely unaffected. By 12 weeks, however, significant reductions in Complex I and IV abundance were evident, accompanied by impaired Complex I activity. Thus, mitochondrial dysfunction develops between 4 and 12 weeks of sustained α-synuclein expression rather than occurring as an immediate consequence of α-synuclein accumulation. Consistent with this progression, the relationship between intraneuronal α-synuclein burden and Complex I abundance was also time-dependent. No significant correlation was detected at 4 weeks, whereas a significant negative correlation emerged at 12 weeks.

A key finding of this study is that this temporal development of mitochondrial dysfunction was paralleled by increasing proximity of α-synuclein to mitochondrial proteins. At 4 weeks, α-synuclein–mitochondrial proximity was relatively limited, whereas by 12 weeks both total and phosphorylated α-synuclein showed significantly increased proximity to mitochondrial proteins, including TOM20, GRP75, and LONP1. Previous studies have reported associations between α-synuclein and mitochondria and consequent disruption of mitochondrial function; however, our findings demonstrate that α-synuclein-mitochondrial proximity increases over the same 4-to-12-week period in which mitochondrial respiratory deficits emerge in nigral dopaminergic neurons in vivo. This temporal relationship raises the possibility that the progressive recruitment or localization of α-synuclein to mitochondria contributes to the development of impaired oxidative phosphorylation.

Another key finding of this study emerged from unbiased spatial transcriptomic analysis, which identified circadian regulation alongside mitochondrial and metabolic processes among the pathways altered following α-synuclein overexpression. Examination of individual circadian-associated genes revealed differential regulation of multiple components of the molecular clock. These included increased expression of *Nr1d1, Nr1d2, Cry2, Arntl2, Csnk1d,* and *Csnk1e*, alongside decreased expression of circadian-associated regulators including *Rorb, Bhlhe41, Rora,* and *Arnt*. Therefore, rather than indicating uniform activation or suppression of the molecular clock, the transcriptomic profile suggests broader remodelling of circadian regulatory pathways in the substantia nigra.

Circadian disturbances are increasingly recognized as a prominent feature of Parkinson’s disease, with patients exhibiting alterations in sleep–wake cycles, hormonal rhythms, and daily activity patterns^5–7^. While these abnormalities are generally considered at the systemic level, circadian clock components also function within individual cells, where they regulate processes extending beyond the generation of organism-level rhythms. These include mitochondrial metabolism, redox homeostasis, oxidative stress responses, and cellular energy production^23,24^. Our findings therefore suggest that, in addition to the systemic circadian abnormalities described in PD, α-synuclein pathology is associated with altered regulation of molecular clock components within vulnerable nigral dopaminergic neurons themselves.

Particularly interesting was the altered regulation of the cryptochrome arm of the molecular clock. The transcriptomic analysis identified increased *Cry2* expression, prompting us to examine CRY1 and CRY2 at the protein level. This revealed a differential shift in cryptochrome expression, characterized by increased CRY2 together with reduced CRY1 immunoreactivity and altered CRY1 distribution. As CRY1 and CRY2 are closely related components of the circadian repressive machinery that are normally coordinated within the molecular clock, their divergent regulation suggests selective remodelling of cryptochrome-dependent regulation rather than a uniform change in clock activity. Importantly, these changes do not by themselves demonstrate altered circadian rhythmicity. Instead, they identify dysregulation of molecular clock components within α-synuclein-expressing dopaminergic neurons, which may have consequences for the intracellular metabolic and stress-response pathways regulated by these proteins.

The relationship between circadian regulation and mitochondrial function is increasingly recognized, with molecular clock components influencing mitochondrial metabolism, oxidative phosphorylation, redox homeostasis, and cellular energy balance^25^. Conversely, mitochondrial metabolic state and redox signaling can influence the molecular clock, indicating that these systems are interconnected rather than operating independently^26^. In this context, the parallel alterations in mitochondrial function and circadian regulatory pathways observed in this study raise the possibility that these processes are linked during α-synuclein-mediated neurodegeneration. Sustained α-synuclein pathology could potentially disrupt the normal coordination between circadian regulation and mitochondrial metabolism, potentially reducing the ability of dopaminergic neurons to adapt their energy production and stress responses to changing cellular demands. In turn, mitochondrial dysfunction, altered redox state, and metabolic stress could further perturb clock-regulated processes, establishing a potentially self-reinforcing cycle of circadian-metabolic dysregulation and neuronal stress. Such a relationship may be particularly relevant to nigral dopaminergic neurons, which have substantial energetic demands and a well-established vulnerability to mitochondrial and oxidative stress.

This potential bidirectional relationship also raises therapeutic possibilities. If disruption of circadian-metabolic coordination contributes to neuronal vulnerability, interventions that restore or stabilize circadian regulation could potentially benefit mitochondrial metabolism, cellular stress responses, and neuronal resilience. Conversely, improving mitochondrial function may help preserve processes required for normal circadian regulation. This possibility is particularly intriguing given evidence that circadian disturbances can occur early in Parkinson’s disease and may therefore represent a modifiable component of disease-associated dysfunction. However, although our data demonstrate molecular and temporal associations between these pathways, they do not establish a causal or bidirectional relationship. Manipulation of individual clock components, including CRY1 and CRY2, will be required to determine whether these alterations contribute to mitochondrial dysfunction or instead represent a parallel or compensatory response to α-synuclein toxicity.

Of note, the transcriptomic analysis was performed at a single time point, and circadian gene expression exhibits marked time-of-day variation. Consequently, the observed transcriptional changes cannot distinguish alterations in circadian phase or amplitude from changes in overall expression. Similarly, measurements of CRY1, CRY2, and BMAL1 at individual experimental time points do not establish altered circadian rhythmicity. Rather, our data demonstrate altered abundance and regulation of molecular clock components within α-synuclein-expressing dopaminergic neurons. Functional assessment across the circadian cycle, together with manipulation of specific clock components, will therefore be important for determining the functional significance of these changes.

In summary, our findings demonstrate that human α-synuclein overexpression in the substantia nigra is associated with progressive mitochondrial dysfunction and selective alterations in circadian regulatory pathways, including differential regulation of the cryptochrome proteins CRY1 and CRY2. Importantly, these findings place two processes increasingly implicated in Parkinson’s disease, mitochondrial dysfunction and circadian dysregulation, within the same vulnerable dopaminergic neuronal population. Rather than establishing a causal relationship between them, our findings raise the possibility that disruption of normal circadian-metabolic coordination represents an underappreciated component of α-synuclein pathology. Determining whether restoration of circadian regulation can preserve mitochondrial function and enhance dopaminergic neuron resilience will be an important direction for future studies.

## Supplemental figures

**Supplemental Figure 1:**
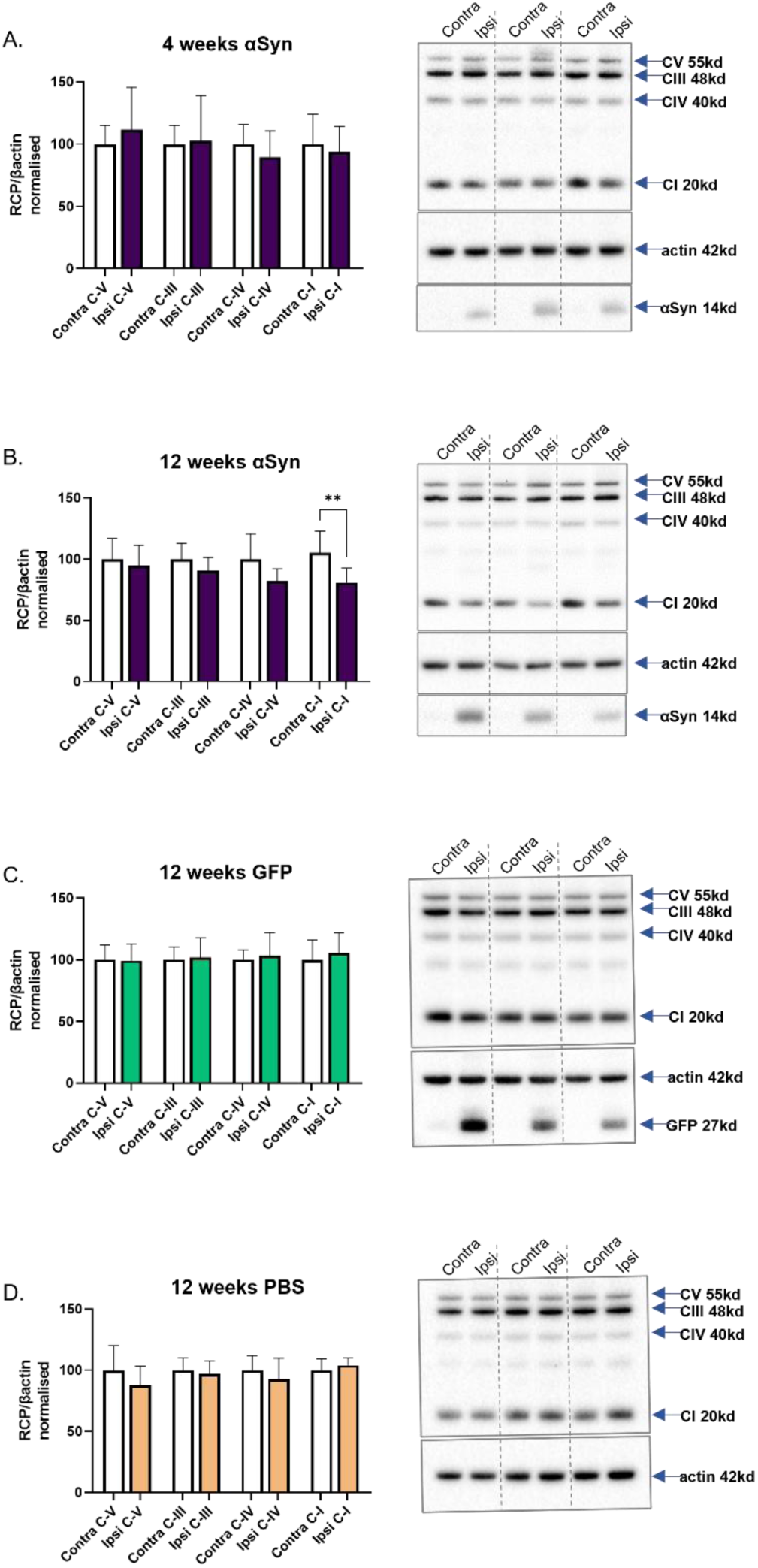
Western blot analysis of mitochondrial respiratory chain proteins following AAV-human α-synuclein, AAV-GFP, or PBS injection. A. Quantification of mitochondrial respiratory chain proteins (RCP) normalized to β-actin at 4 weeks following AAV-human α-synuclein injection, with representative immunoblots of contralateral and ipsilateral substantia nigra shown beside. B. Quantification and representative immunoblots at 12 weeks following AAV-human α-synuclein injection. C. Quantification and representative immunoblots from AAV-GFP-injected mice at 12 weeks. D. Quantification and representative immunoblots from PBS-injected mice at 12 weeks.

**Supplemental Figure 2:**
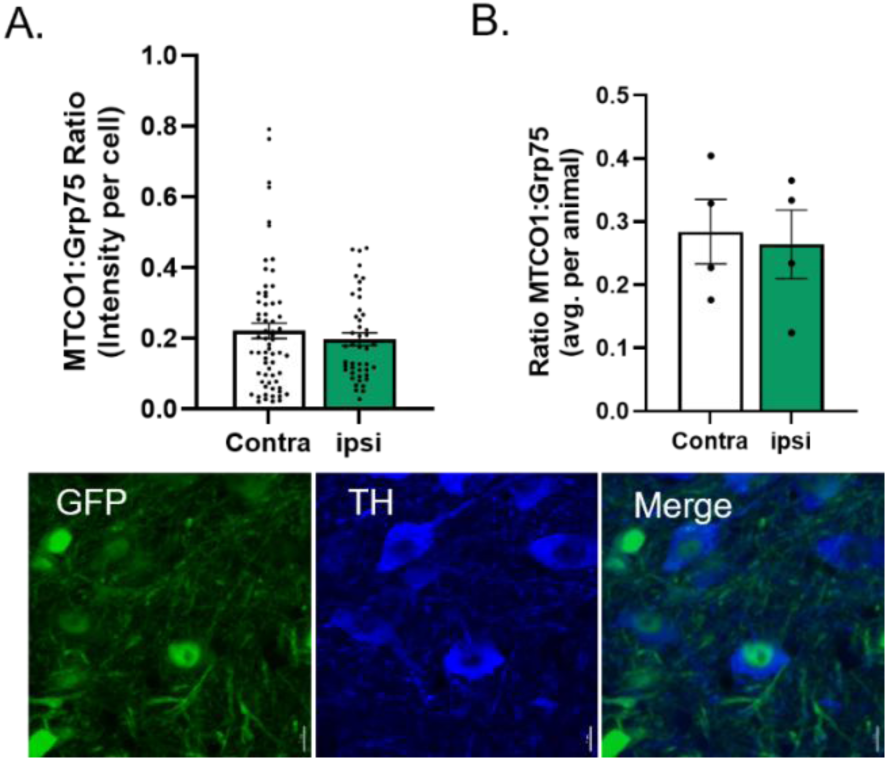
AAV-GFP injection does not alter complex IV protein expression in dopaminergic neurons. A. Per-cell immunofluorescent quantification of the MTCO1:GRP75 ratio (Complex IV) in TH-positive neurons of AAV-GFP-injected mice. B. Per-animal quantification of the MTCO1:GRP75 ratio, with representative images showing GFP expression in substantia nigra neurons.

**Supplemental Figure 3:**
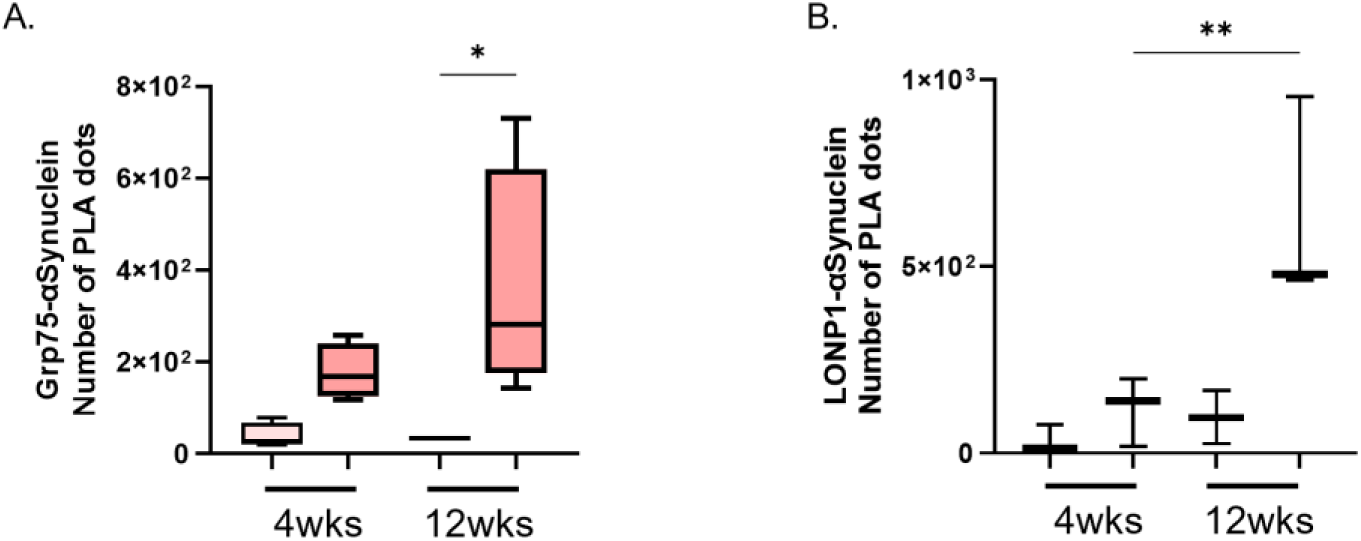
α-Synuclein association with mitochondrial proteins increases over time in the substantia nigra. Positive PLA puncta were quantified within the substantia nigra pars compacta (SNpc). A,B. Quantification and representative images of TOM20–Syn211 PLA signal at 4 and 12 weeks following α-synuclein overexpression. Graphs show quantification of PLA in contra- and ipsilateral SNpc of mice overexpressing human αSyn. C-D. Quantification of TOM20–pS129, GRP75–pS129 PLA signal. N=4-5 mice per group, statistics used were student’s t-test and one-way ANOVA. *p<0.05, **p<0.01.

